# Characterization and mu opioid receptor sensitivity of neuropeptide Y interneurons in the nucleus accumbens

**DOI:** 10.1101/2022.05.02.490329

**Authors:** Cassandra L. Retzlaff, Patrick E. Rothwell

## Abstract

Inhibitory interneurons represent less than 5% of neurons within the nucleus accumbens, but are critical for proper microcircuit function within this brain region. In the dorsal striatum, neuropeptide Y is expressed by two interneuron subtypes (low-threshold spiking interneurons and neurogliaform interneurons) that exhibit mu opioid receptor sensitivity in other brain regions. However, few studies have assessed the molecular and physiological properties of neuropeptide Y interneurons within the nucleus accumbens. We used a transgenic reporter mouse to identify and characterize neuropeptide Y interneurons in acute nucleus accumbens brain slices. Nearly all cells exhibited electrophysiological properties of low-threshold spiking interneurons, with almost no neurogliaform interneurons observed among neuropeptide Y interneurons. We corroborated this pattern using fluorescent in situ hybridization, and also identified a high level of mu opioid receptor expression by low-threshold spiking interneurons, which led us to examine the functional consequences of mu opioid receptor activation in these cells using electrophysiology. Mu opioid receptor activation caused a reduction in the rate of spontaneous action potentials in low-threshold spiking interneurons, as well as a decrease in optogenetically-evoked GABA release onto medium spiny neurons. The latter effect was more robust in female versus male mice, and when the postsynaptic medium spiny neuron expressed the Drd1 dopamine receptor. This work is the first to examine the physiological properties of neuropeptide Y interneurons in the nucleus accumbens, and show they may be an important target for mu opioid receptor modulation by endogenous and exogenous opioids.

## 1. INTRODUCTION

The nucleus accumbens is comprised of mostly GABAergic neurons, including a large number of medium spiny neurons (MSNs) that project to other brain structures, as well as a small number of inhibitory interneurons that form connections within local microcircuitry (Burke et al., 2017; Castro and Bruchas, 2019). While inhibitory interneurons represent <5% of all neurons in the nucleus accumbens, they have important roles in regulating MSN output through direct monosynaptic inhibition (Gittis et al., 2010; Koos and Tepper, 1999; Scudder et al., 2018; Straub et al., 2016), as well as more complex synaptic mechanisms (English et al., 2011; Fino et al., 2018; Holly et al., 2021). Different subtypes of inhibitory interneurons can be defined on the basis of their electrophysiological properties and molecular markers (Kawaguchi, 1993). For example, fast-spiking interneurons that express parvalbumin are well-studied and represent a major source of feed-forward inhibition within the nucleus accumbens and dorsal striatum (Owen et al., 2018; Scudder et al., 2018; Straub et al., 2016; Yu et al., 2017).

Neuropeptide Y (NPY) is a marker expressed by two other inhibitory neuron subtypes in the striatum: low-threshold spiking interneurons (LTSIs) and neurogliaform interneurons (Ibanez-Sandoval et al., 2011). LTSIs co-express NPY as well as other unique molecular markers like somatostatin (SST) and nitric oxide synthase, whereas neurogliaform interneurons express NPY but not SST (Ibanez-Sandoval et al., 2011; Kawaguchi, 1993). Both NPY interneuron subtypes have been studied in the dorsal striatum (English et al., 2011; Holly et al., 2019; Holly et al., 2021), but less in known about the properties and function of LTSIs in the nucleus accumbens (Ribeiro et al., 2019; Ribeiro et al., 2018; Scudder et al., 2018), while neurogliaform interneurons in the nucleus accumbens have not previously been characterized.

Given the key role of the nucleus accumbens in reward and addiction, it is notable that LTSIs and neurogliaform interneurons exhibit mu opioid receptor (MOR) sensitivity in other brain regions (Elghaba and Bracci, 2017; Krook-Magnuson et al., 2011). In this study, we conduct a comprehensive characterization of NPY interneurons in the nucleus accumbens, including their MOR expression and functional sensitivity to MOR agonists. We find that nearly all NPY interneurons in the nucleus accumbens have the physiological and molecular profile of LTSIs, and MOR activation inhibits their intrinsic excitability as well as their release of GABA onto MSNs. These results show that NPY interneurons represent an unappreciated target for opioid signalling within the nucleus accumbens, and may contribute to the behavioral consequences of MOR activation within this brain region.

## 2. MATERIALS AND METHODS

### 2.1 Subjects

NPY-GFP reporter mice (van den Pol et al., 2009) were obtained from The Jackson Laboratory (strain #006417), and maintained by breeding hemizygotes with C57Bl/6J partners. Sst-ires-Flp mice (He et al., 2016) were obtained from The Jackson Laboratory (strain #031629), and crossed with Drd1-tdTomato/Drd2-eGFP double-transgenic reporter mice (Trieu et al., 2022) on a C57Bl6J genetic background. Mice were housed in groups of 2-5 per cage, on a 12 hour light cycle (0600h – 1800h) at ∼23° C with food and water provided ad libitum. Experimental procedures were conducted between 1000h – 1600h and were approved by the Institutional Animal Care and Use Committee of the University of Minnesota.

### 2.2 Fluorescent *in situ* hybridization

C57Bl/6J wild-type mice were anesthetized with isoflurane and brains were rapidly dissected and frozen on dry ice in OCT embedding medium and stored at -80°C until being sectioned on a cryostat (Leica CM3050s). Ten-micron sections were collected at -20°C, placed immediately on positively charged glass slides, and stored at -80°C until ready for RNAscope processing. Protocols and reagents were provided by Advanced Cell Diagnostics (ACD) and used to conduct the RNAscope assay as previously described (Pisansky et al., 2019). Briefly, slides were post-fixed in 4% PFA for 15 min at 4°C followed by ethanol dehydration. Slides were dehydrated in increasing concentrations of ethanol (50%>70%>100%) for 5 min each at room temperature. After final 100% ethanol wash, slides were stored at -20°C in 100% ethanol overnight or used immediately for processing. Slides were then left to air dry, and a hydrophobic barrier was painted around sections that would be used for mRNA detection. Protease IV was applied to sections and let to sit for 30 min at room temperature. Slides were washed quickly twice in PBS. Following the PBS washes, hybridizing probes (Sst, Npy, and Oprm1) were applied to sections and incubated at 40°C for 2 hours, and then washed twice in 1x RNAscope buffer for 2 min. The following steps were a series of amplification steps with provided AMP1-4 detection reagents. The reagents were applied sequentially and incubated for 15-30 min each at 40°C as per the instruction manual and washed with 1X RNAscope between each reagent. After the final wash, slides were dried and mounting medium (Invitrogen Prolong Gold antifade reagent with DAPI) was applied before coverslips were placed on the slides and stored at 4°C. Images were taken within the following week on a Keyence BZX fluorescence microscope under a 10 or 40x objective.

### 2.3 Stereotaxic Surgery

To enable optogenetic stimulation of LTSI synapses onto MSNs, we performed stereotaxic injection of adeno-associated virus (AAV) into the nucleus accumbens, as previously described (Pisansky et al., 2019). Briefly, Sst-ires-Flp mice were anesthetized with a ketamine:xylazine cocktail (100:10 mg/kg), and a small hole was drilled above target coordinates for the nucleus accumbens (AP +1.5, ML +0.8; DV -4.6). The viral solution injected into the brain contained two AAV vectors in an equal mixture:

- AAVdj-CAG-fDIO-GFP-Cre (Penzo et al., 2015)
- AAVdj-EF1a-DIO-ChIEF-Venus (Pisansky et al., 2019)

A nanoject syringe affixed with a glass pipette containing viral solution was slowly lowered to these target coordinates, followed by injection of 0.5 μL at a rate of 0.1 μL/min. The syringe tip was left in place at the injection site for 5 minutes, and then slowly retracted over the course of 5 minutes. Electrophysiological experiments were conducted 4-6 weeks after surgery, to allow for robust virus expression.

### 2.4 Electrophysiology

As previously described (Toddes et al., 2021), mice were anesthetized with isoflurane and perfused with ice cold sucrose solution containing (in mM): 228 sucrose, 26 NaHCO3, 11 glucose, 2.5 KCl, 1 NaH2PO4-H2O, 7 MgSO4-7H20, 0.5 CaCl2-2H2O. Brains were then rapidly dissected and placed in cold sucrose. Coronal slices (240 μm thick) containing the nucleus accumbens were collected using a vibratome (Leica VT1000S) and allowed to recover in a submerged holding chamber with artificial cerebrospinal fluid (aCSF) containing (in mM): 119 NaCl, 26.2 NaHCO3, 2.5 KCl, 1 NaH2PO4-H2O, 11 glucose, 1.3 MgSO4-7H2O, 2.5 CaCl2-2H2O. Slices recovered in warm ACSF (33°C) for 10-15 minutes and then equilibrated to room temperature for at least one hour before use. Slices were transferred to a submerged recording chamber and continuously perfused with aCSF at a rate of 2 mL/min at room temperature. All solutions were continuously oxygenated (95% O_2_/5% CO_2_).

Current clamp recordings were conducted using the NPY-GFP reporter line to identify LTSI or neurogliaform interneurons within the nucleus accumbens or dorsal striatum. Recording electrodes made of borosilicate glass (3-5 MΩ) were filled with (in mM): 120 K-Gluconate, 20 KCl, 10 HEPES, 0.2 EGTA, 2 MgCl2, 4 ATP-Mg, 0.3 GTP-Na (pH 7.2-7.3). Cells were injected with a series of current steps (0.5 sec duration) from - 100 to +450 pA. For cell-attached recordings from LTSIs, cells were identified by GFP expressed under the NPY promoter, and by the presence of spontaneous action potentials. Recording electrodes were made of borosilicate glass electrodes (1-2 MΩ) and filled with aCSF. Bath aCSF contained atropine (10 µM) and mecamylamine (20 µM) to block cholinergic transmission, followed by wash on of picrotoxin (50 µM), NBQX (10 µM), and D-APV (50 µM) to block GABA_A_ and glutamate receptors during measurement of baseline spontaneous activity. Seals on the cell membrane were achieved through negative pressure, but membranes were never broken. Seal resistances were held from 20 mOhm to 1 gOhm, and action potential spikes were included for analysis if over a 15 pA cutoff. Cells were only included in the analysis if the baseline firing rate was stable prior to DAMGO wash.

For optogenetic stimulation of GABA release onto MSNs, whole-cell voltage clamp recordings were obtained under visual control using DIC optics. D1- and D2-MSNs were distinguished by the presence of tdTomato or GFP fluorescence, respectively, using an Olympus BX51W1 microscope. In instances where SST-Flp/D2-GFP animals were injected with viruses to express ChIEF, which was also fused to a green fluorophore (Venus), cells were distinguished by their response to optogenetic stimulation. Cells that expressed the opsin exhibited a very large and uncharacteristic current and were excluded from further analysis. Voltage clamp recording electrodes were made of borosilicate glass electrodes (3-5 MΩ) and filled with (in mM): 125 CsCl, 10 TEA-Cl, 10 HEPES, 0.1 EGTA, 3.3 QX-314 (Cl− salt), 1.8 MgCl2, 4 Na2-ATP, 0.3 Na-GTP, 8 Na2-phosphocreatine (pH 7.3 adjusted with CsOH; 275-280 mOsm). Extracellular bath solution contained NBQX (10 µM), D-APV (50 µM), atropine (10 µM), and mecamylamine (20 µM). Optically-evoked inhibitory postsynaptic currents (oIPSCs) were driven by 475 nm light pulses (1 mW power) with a 2-5 ms pulse width, while holding the postsynaptic MSN at -70 mV. All recordings were performed using a MultiClamp 700B amplifier (Molecular Devices), filtered at 2 kHz, and digitized at 10 kHz. Data acquisition and analysis were performed online using Axograph software. Series resistance, when applicable, was monitored continuously and experiments were discarded if resistance changed by >20%.

### 2.5 Statistical Analyses

Similar numbers of male and female animals were used in all experiments, with samples size indicated in figure legends. Individual data points from males (filled circles) and females (open circles) are distinguished in each figure. Sex was included as a variable in factorial ANOVA models analyzed using GraphPad Prism software, with repeated measures on within-cell factors. Sex differences were not statistically significant unless noted otherwise, and in the absence of a sex difference, simple main effects were further analyzed using t-tests. The Type I error rate was set to α = 0.05 (two-tailed) for all comparisons. All summary data are displayed as mean ± SEM.

## 3. RESULTS

### 3.1 Electrophysiological profile of NPY interneurons in the nucleus accumbens and dorsal striatum

To characterize the electrophysiological properties of nucleus accumbens interneurons that express NPY, we prepared acute coronal brain slices from NPY-GFP transgenic reporter mice (van den Pol et al., 2009) (Fig. 1A). We conducted whole-cell current-clamp recordings from green neurons in the nucleus accumbens and identified two subtypes of cells with distinct electrophysiological properties, as previously reported in the dorsal striatum (Ibanez-Sandoval et al., 2011). The first subtype had a depolarized resting membrane potential, high input resistance, and low threshold to fire action potentials (Fig. 1B). These properties match prior descriptions of LTSIs in the nucleus accumbens (Ribeiro et al., 2018; Scudder et al., 2018) and dorsal striatum (Kawaguchi, 1993). The second subtype had a hyperpolarized resting membrane potential, low input resistance, and high threshold to fire action potentials (Fig. 1C). These properties match prior descriptions of neurogliaform interneurons in the dorsal striatum (Ibanez-Sandoval et al., 2011).

**Figure 1.**
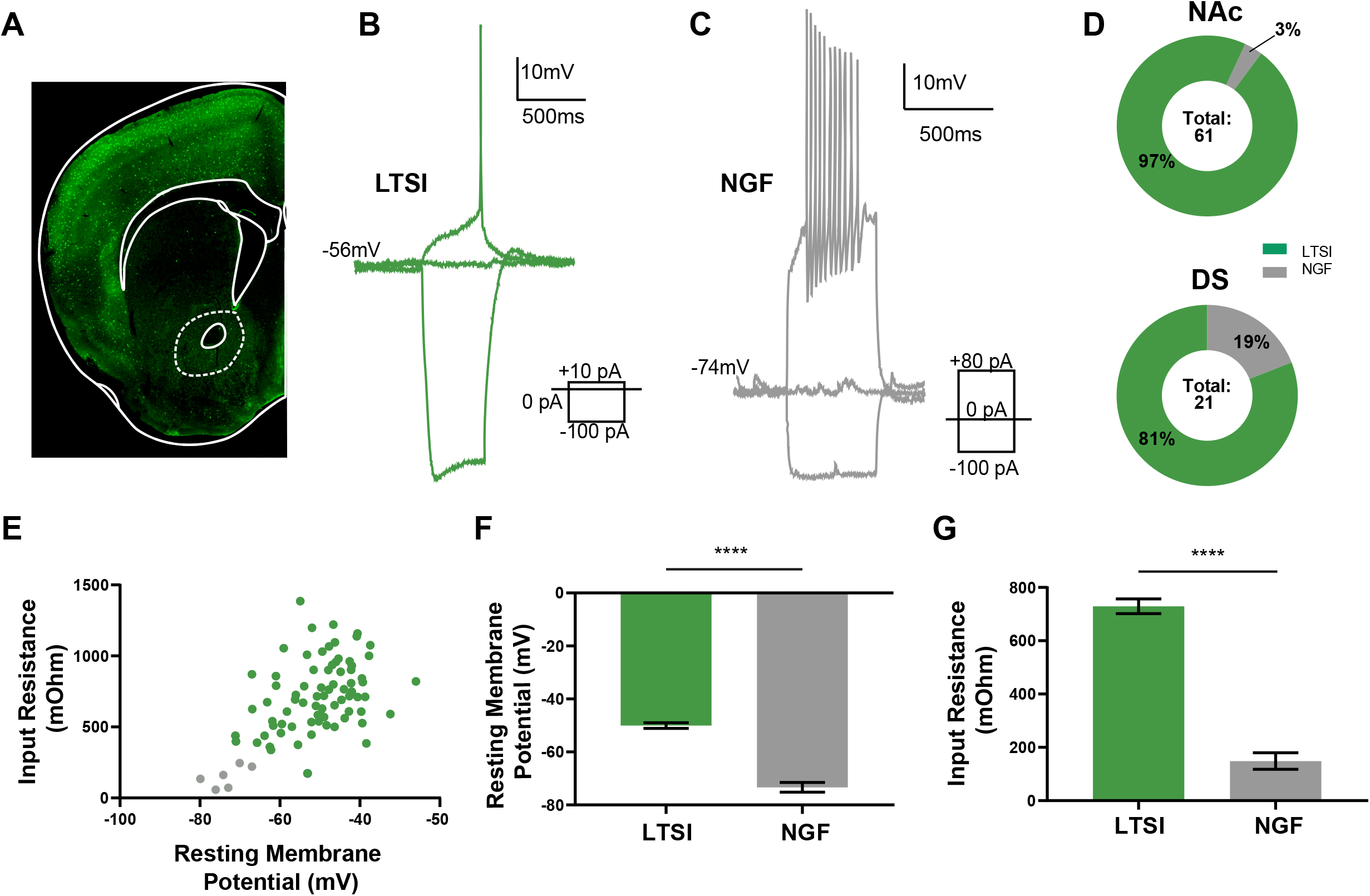
Nucleus accumbens NPY neurons are predominately LTSIs. A. Coronal brain section from an NPY-GFP transgenic mouse used to identify NPY interneurons for electrophysiological characterization. B. Whole-cell current clamp-recording from an NPY-GFP neuron with LTSI properties. C. Whole-cell current-clamp recording from an NPY-GFP neuron with neurogliaform properties. D. Percentage of NPY-GFP neurons with LTSI or neurogliaform properties recorded in the nucleus accumbens (NAc) or dorsal striatum (DS). E. Scatter plot showing the resting membrane potential and input resistance of individual NPY-GFP neurons characterized as LTSI (green) or neurogliaform (gray). F. Average resting membrane potential for LTSI and neurogliaform-like neurons. G. Average input resistance for LTSI and neurogliaform-like neurons. **** indicates p < 0.0001.

In the nucleus accumbens, the vast majority of cells expressing GFP (59/61) had electrophysiological properties of LTSIs, and only a very small number (2/61) had electrophysiological properties of neurogliaform interneurons (Fig. 1D). The relative proportion of each cell type among GFP-positive neurons in the nucleus accumbens (∼97% LTSI, ∼3% neurogliaform) was somewhat different than a prior characterization of GFP-positive neurons in the dorsal striatum of this same mouse line (∼79% LTSI, ∼21% neurogliaform) (Ibanez-Sandoval et al., 2011). To cross-validate our analysis in the nucleus accumbens, we also recorded 21 cells expressing GFP in the dorsal striatum. Seventeen of these cells had electrophysiological properties of LTSIs, and the remaining four cells had electrophysiological properties of neurogliaform interneurons (Fig. 1D). The relative proportion of each cell type among GFP-positive neurons in the dorsal striatum (∼81% LTSI, ∼19% neurogliaform) was thus quite similar to prior reports (Ibanez-Sandoval et al., 2011). An aggregate analysis of all cells expressing GFP in both the nucleus accumbens and dorsal striatum clearly illustrates the distinct electrophysiological properties of LTSIs and neurogliaform interneurons (Fig. 1E-G).

### 3.2 Molecular profile and MOR expression of NPY neurons in the nucleus accumbens

We were surprised that so many NPY-GFP cells in the nucleus accumbens were LTSIs, and so few were neurogliaform interneurons. To further investigate the molecular profile of NPY neurons in nucleus accumbens tissue from wild-type C57Bl/6J mice, we used the RNAscope platform to perform fluorescent in situ hybridization (Fig. 2A). Previous reports indicate neurons with LTSI electrophysiological properties show co-localized expression of NPY with SST (Ibanez-Sandoval et al., 2011), so we simultaneously stained tissue with probes for both NPY and SST mRNA, as well as a third probe for MOR mRNA (Fig. 2B). We identified 109 cells in the nucleus accumbens that expressed NPY mRNA, and found that 92% of them (101/109) also expressed SST mRNA (Fig. 2C), corroborating our electrophysiological analysis and confirming the abundance of LTSIs in the nucleus accumbens. The percentage of NPY cells expressing SST was similar in tissue from female mice (95%) and male mice (90%). Cells expressing NPY mRNA were evenly distributed throughout different nucleus accumbens subregions, including the medial shell (27%), ventral shell (34%), and core (37%).

**Figure 2.**
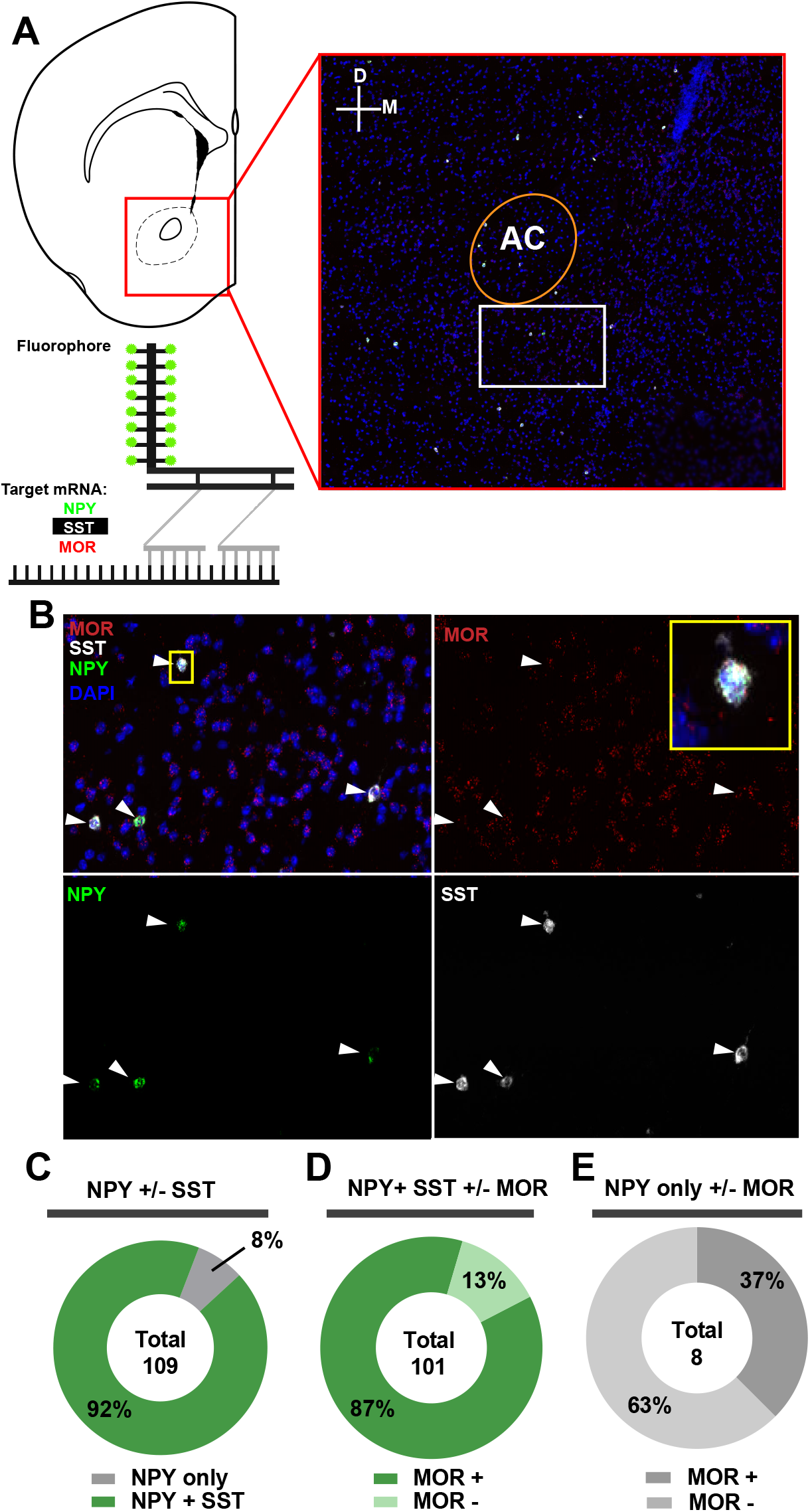
Fluorescent in situ hybridization reveals expression of MOR by LTSIs. A. Schematic showing a coronal brain section from a wild-type mouse, highlighting the nucleus accumbens (red box) and RNAscope probes used to detect NPY (green), SST (white), and MOR (red) at 10X magnification. Schematic of RNAscope platform adapted from ACDbio and created at biorender.com. B. Individual and merged images at 40x showing co-localization of MOR, NPY, and SST mRNA within the region highlighted in panel A (white box). C. Quantification of NPY-expressing cells in the nucleus accumbens that also express SST (92%). D. Quantification of NPY and SST co-expressing cells that also express MOR (87%). E. Quantification of NPY-expressing cells that do not express SST but do express MOR (37%).

We also quantified co-expression of MOR mRNA by NPY cells (Fig. 2D). The majority (87%) of neurons expressing NPY and SST (putative LTSIs) also expressed MOR mRNA. This percentage was high in tissue from male mice (80%) and female mice (95%). Among the small number of nucleus accumbens cells expressing NPY but not SST (putative neurogliaform interneurons), only 30% also expressed MOR mRNA. In light of this convergent evidence for a low abundance of neurogliaform interneurons in the nucleus accumbens, we did not pursue further analysis of this interneuron subtype. Instead, we focused the remainder of this study on LTSIs, given their greater abundance and higher level of MOR expression.

### 3.3 MOR activation reduces LTSI spontaneous activity

To assess the functional effects of MOR activation on LTSI activity, we again used electrophysiology from NPY-GFP mice (Fig. 3A). We leveraged the fact that LTSIs are spontaneously active in acute nucleus accumbens brain slices (Scudder et al., 2018), and recorded in a cell-attached configuration to assess changes in spontaneous activity after MOR activation (Fig. 3B). To study the direct effects of MOR activation on LTSI activity, these recordings were performed in the presence of atropine (10 µM) and mecamylamine (20 µM), to block acetylcholine receptors and mitigate the indirect influence of MOR activation on cholinergic transmission (Elghaba and Bracci, 2017). The recording solution also contained antagonists for AMPA receptors (10 µM NBQX), NMDA receptors (50 µM D-APV), and GABA_A_ receptors (50 µM picrotoxin), in order to avoid any indirect influence of MOR activation on excitatory or inhibitory synaptic input to LTSIs.

**Figure 3.**
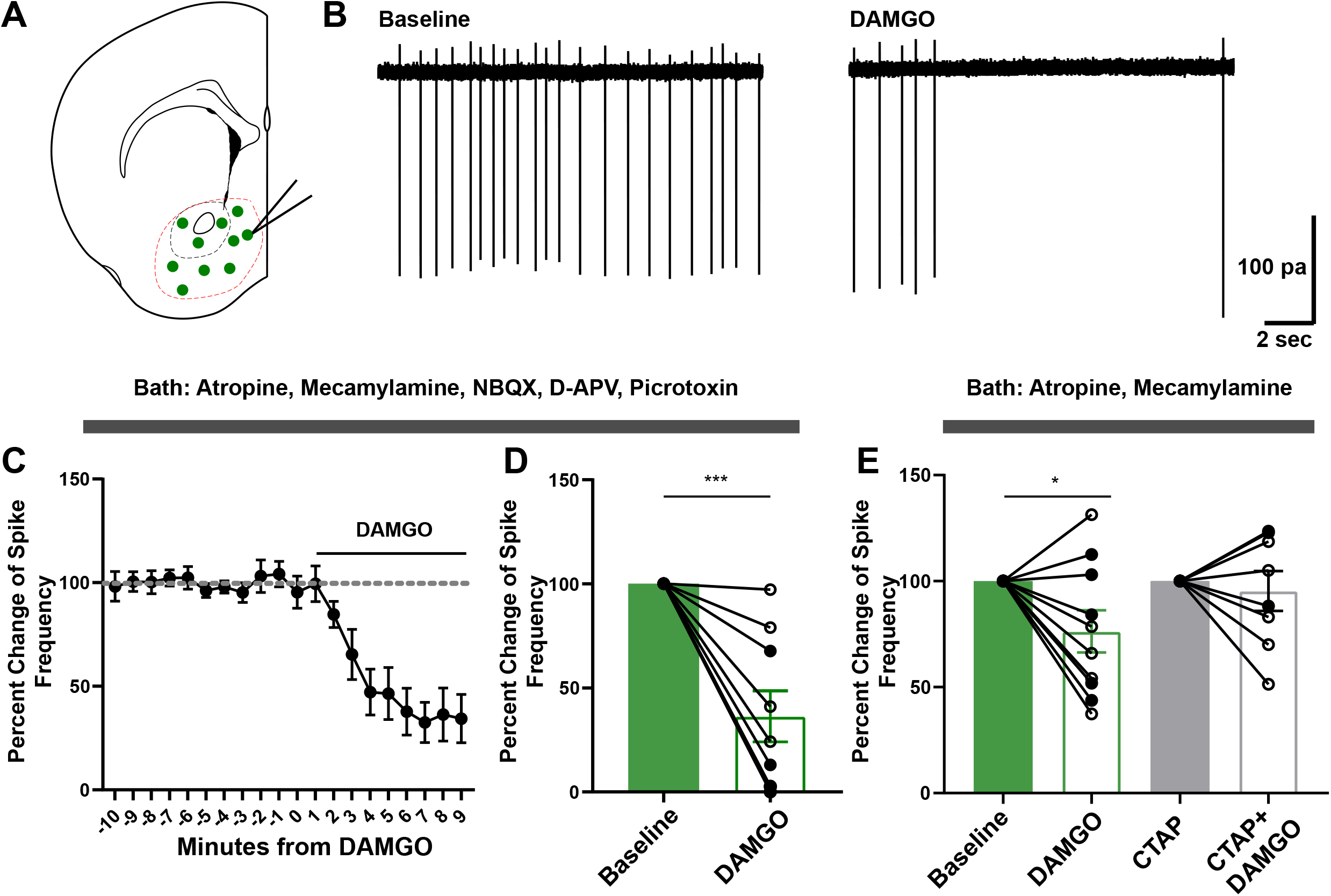
MOR activation reduces the spontaneous activity of LTSIs. A. Schematic showing LTSIs in the nucleus accumbens of an NPY-GFP mouse, which were targeted for cell-attached recording. B. Example traces of spontaneous firing from an LTSI before and after DAMGO (300 nM) application. C. Time course of spontaneous activity during DAMGO bath application in 9 cells (5 female, 4 male) from 5 mice (3 female, 2 male). D) Quantification of percent change from baseline during the final two minutes of recording in the presence of cholinergic, glutamate, and GABAA receptor antagonists. E. Quantification of DAMGO effect in the presence of only cholinergic blockers in 10 cells (5 female, 5 male) from 8 mice (3 female, 5 male), and in the added presence of CTAP (1 µM) in 8 cells (5 female, 3 male) from 7 mice (4 female, 3 male). *** indicates p < 0.001; * indicates p < 0.05.

MOR activation was induced by a 10 minute bath application of DAMGO, at a concentration (300 nM) shown to selectively activate MOR in striatal brain slices (Banghart et al., 2015). DAMGO application caused a striking and significant decrease in the spontaneous firing rate of LTSIs (Fig. 3C), with a reduction of 63.6% compared to baseline (t_8_ = 5.20, p = 0.0008). This decrease in firing rate was observed in cells from both female mice (79.2%) and male mice (51.2%) (Fig. 3D). We also conducted this experiment in the absence of NBQX, D-APV, and picrotoxin. Under these conditions, the decrease in firing rate was smaller (23.7% reduction) but still reliable (t_9_ = 2.38, p = 0.041), and blocked by the MOR antagonist CTAP (1µM) (4.7% reduction in firing rate; t_8_ < 1). These data indicate that MOR activation reduced the spontaneous activity of LTSIs within the nucleus accumbens.

### 3.4 MOR activation decreases GABA output onto MSNs of the nucleus accumbens

To determine if MOR activation also modulated the GABAergic output of LTSIs onto MSNs, we utilized an optogenetic strategy (Scudder et al., 2018; Straub et al., 2016) involving expression of ChIEF, an excitatory opsin (Lin et al., 2009). Sst-ires-Flp mice received stereotaxic co-injection of two AAV vectors into the nucleus accumbens (Fig. 3A). The first expressed Cre recombinase in a Flp-dependent fashion (AAVdj-CAG-fDIO-GFP-Cre), while the second expressed ChIEF in a Cre-dependent fashion (AAVdj-EF1a-DIO-ChIEF-Venus). This dual-recombinase strategy was used to enable forward-compatibility with other Cre-dependent viral and genetic manipulations in future studies. Mice used in experiments also carried Drd1-tdTomato and/or Drd2-eGFP reported transgenes, enabling us to differentiate recordings from D1-MSNs and D2-MSNs.

To confirm functional opsin expression, LTSIs expressing ChIEF-Venus were recorded in voltage-clamp and stimulated with a one-second pulse of blue light. As expected, the LTSI responded with a sharp inward current, followed by a slight decay to a steady-state plateau that lasted throughout the duration of the light pulse (Fig. 4B). Recordings from either D1- or D2-MSNs revealed reliable oIPSCs in both MSN subtypes (Fig 4C-D). After a 10 min bath application of DAMGO (300 nM), we found a significant reduction in the amplitude of oIPSC in both D1-MSNs (Fig. 4C; t_9_ = 7.784, p < 0.0001) and D2-MSNs (Fig. 4D; t_7_ = 3.367, p = 0.012). ANOVA revealed a main effect of Cell Type (F_1,14_ = 7.328, p = 0.017), indicating the effect of DAMGO was greater in D1-MSNs than D2-MSNs. There was also a main effect of Sex (F_1,14_ = 19.86, p = 0.0005), indicating the effect of DAMGO was greater in females than males. These recordings also showed a significant increase in the paired-pulse ratio in D1-MSNs (Fig. 4C; t_9_ = 2.33, p = 0.044) as well as D2-MSNs (Fig. 4D; t_7_ = 3.71, p = 0.0075), consistent with a decrease in the probability of GABA release from the presynaptic terminal caused by MOR activation.

**Figure 4.**
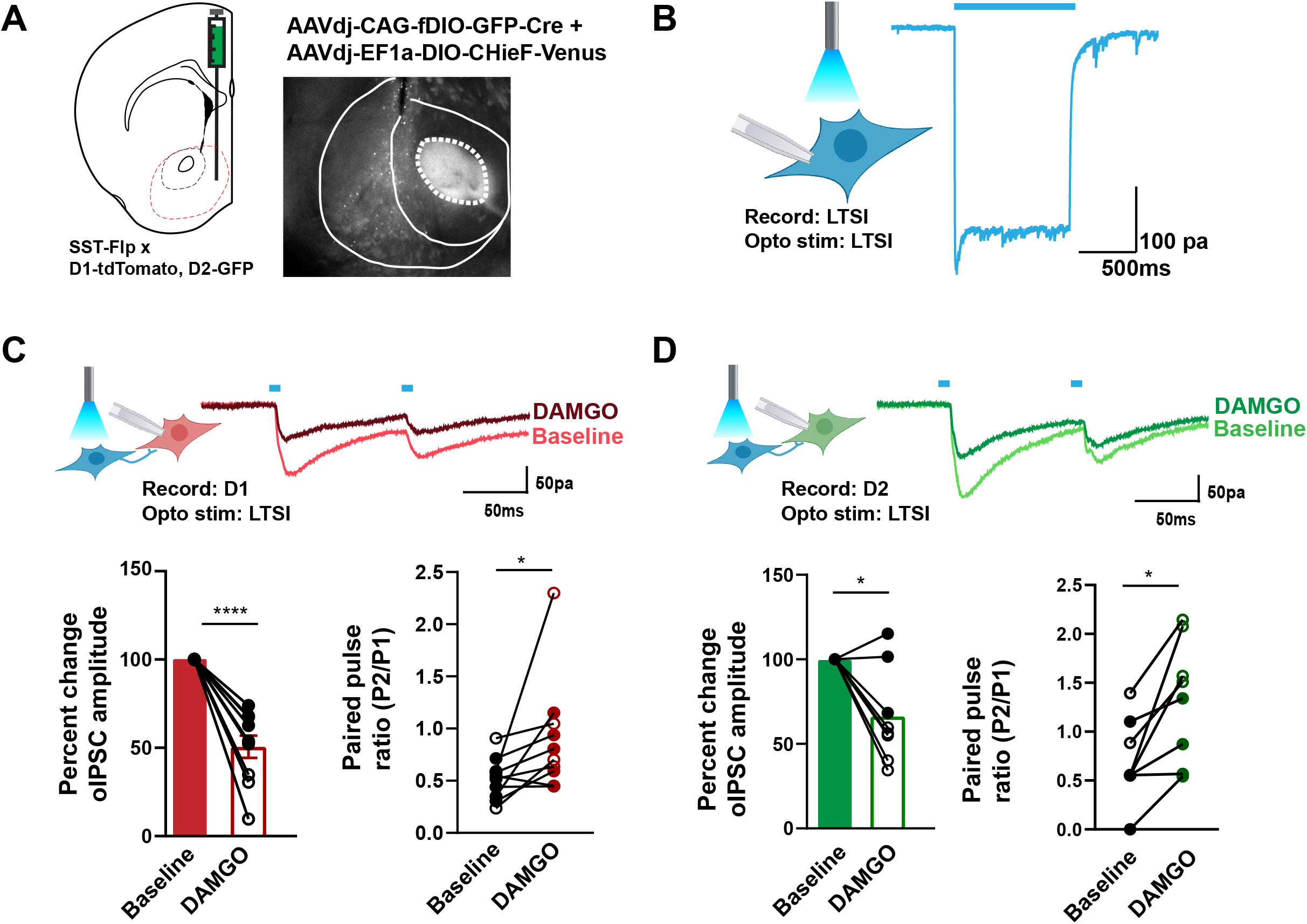
MOR activation decreases GABA release from LTSIs onto MSNs. A. Schematic showing stereotaxic co-injection of AAVdj-CAG-fDIO-GFP-Cre and AAVdj-EF1a-DIO-ChIEF-Venus into the nucleus accumbens of Sst-ires-Flp mice carrying Drd1-tdTomato and/or Drd2-eGFP transgenes. B. Voltage-clamp recording from an LTSI expressing ChIEF, showing the response to one second of optical stimulation (473 nm). C. Top: whole-cell voltage-clamp recording from a D1-MSN, showing example traces of oIPSC recorded before and after DAMGO application. Bottom: quantification of oIPSC amplitude (left) and paired-pulse ratio (right) recorded from 10 D1-MSNs (6 female, 4 male) in 9 mice (5 female, 4 male), before and after 10-min application of DAMGO (300 nM). D. Top: whole-cell voltage-clamp recordings from a D2-MSN, showing example traces of oIPSC recorded before and after DAMGO application. Bottom: quantification of oIPSC amplitude (left) and paired-pulse ratio (right) recorded from 8 D2-MSNs (5 female, 3 male) in 8 mice (5 female, 3 male), before and after 10-min application of DAMGO (300 nM). **** indicates p < 0.0001; * indicates p < 0.05.

## 4. DISCUSSION

This study is the first to characterize NPY interneurons in the nucleus accumbens, including MOR expression and function. We find that nearly all nucleus accumbens NPY interneurons exhibit LTSI properties, as well as a high degree of MOR sensitivity. Neurogliaform interneurons, which represent ∼20% of NPY interneurons in the dorsal striatum (Ibanez-Sandoval et al., 2011), appear to be much less abundant though not entirely absent in the nucleus accumbens. We initially observed this pattern using NPY-GFP reporter mice, and then corroborated it in the nucleus accumbens of wildtype mice using fluorescent in situ hybridization. This gradient in the prevalence of neurogliaform interneurons is similar to the fast-spiking interneurons, which are also more abundant in the dorsal striatum than in the nucleus accumbens (Berke et al., 2004; Kita et al., 1990). Together, these data indicate fundamental differences in the organization of inhibitory microcircuits within dorsal and ventral striatal subregions and show that LTSIs are a prominent feature of nucleus accumbens microcircuitry.

Our fluorescent in situ hybridization analysis also showed a high degree of MOR co-expression by LTSIs, hinting at a functional sensitivity to MOR agonists, as reported for NPY interneurons in other brain regions (Elghaba and Bracci, 2017; Krook-Magnuson et al., 2011). We first evaluated this possibility using cell-attached recordings in acute brain slices to measure the firing rate of spontaneous action potentials. DAMGO application caused a substantial decrease in spontaneous activity, which was most robust in the presence of glutamate and GABA receptor antagonists. Prior work has also shown that the baseline firing rate of LTSIs is increased when GABA_A_ receptors are blocked (Scudder et al., 2018). Together, these results suggest inhibitory inputs to LTSIs play an important role in modulating firing rate, and these inhibitory inputs may also exhibit MOR sensitivity.

Using an optogenetic strategy to stimulate LTSI synapses onto MSNs (Scudder et al., 2018; Straub et al., 2016), we also found that DAMGO decreased the release of GABA from LTSI axon terminals, consistent with presynaptic MOR activation. An interesting and unexpected finding was that this effect of DAMGO was larger in tissue from female mice versus male mice. In our fluorescent in situ hybridization analysis, the co-localized expression of MOR mRNA was more commonly observed in LTSIs from female mice (95%) versus male mice (80%). This difference in the fraction of LTSIs expressing MOR, or a quantitative difference in the amount of MOR expression by individual LTSIs, could explain why female mice more robustly respond to exogenous MOR agonists in our electrophysiological studies. This finding adds to a growing list of sex differences in MOR signaling that may differentially influence behavioral responses to exogenous opioids and susceptibility to opioid use disorders (Becker and Chartoff, 2019).

In addition to a sex difference in MOR inhibition of LTSI GABA release onto MSNs, we also found that the magnitude of this effect depended on the identity of the postsynaptic MSN. GABA release onto D1-MSNs was attenuated by DAMGO to a greater extent than GABA release onto D2-MSNs. Previous studies have shown the oIPSC amplitude and kinetics produced by optogenetic stimulation of LTSIs is similar in D1-MSNs and D2-MSNs (Scudder et al., 2018; Straub et al., 2016), indicating that the difference we observe between MSN subtypes is specifically related to modulation of GABA release by MOR activation, rather than a difference in basal synaptic strength or connectivity. This target-specific effect on LTSI output to MSN subtypes is reminiscent of the target-specific remodeling of fast-spiking interneuron output that occurs after dopamine depletion (Gittis et al., 2011). Given the target-specific MOR sensitivity of LTSI output to MSNs, it will be interesting for future studies to determine if chronic opioid exposure reorganizes LTSI output to MSNs in a target-specific manner.

In this regard, it is worth noting that the MOR sensitivity of LTSIs has important implications for how these interneurons may contribute to natural and pathological states of motivation. The target-specific regulation of GABA release onto MSNs means that local MOR activation would disinhibit D1-MSNs to a greater degree than D2-MSNs. The resulting disparity in MSN output (favoring D1-MSNs) could promote naturally rewarding behaviors like food consumption (Bakshi and Kelley, 1993; Castro et al., 2021) or social interaction (Gunaydin et al., 2014; Trezza et al., 2011), as well as the rewarding properties of exogenous opioids (Koo et al., 2014; O’Neal et al., 2022). It is also fascinating to note that LTSIs in the dorsal striatum exert an inhibitory influence on dopamine release (Holly et al., 2021). If a similar mechanism exists in the nucleus accumbens, then MOR inhibition of LTSI output could enhance motivation via disinhibition of dopamine release. These hypotheses can be tested in future studies by using Sst-ires-Flp mice and Flp-dependent Cre virus to perform localized cell type-specific genetic knockout of MOR expression – a manipulation successfully used in the dorsal striatum to manipulate GABA release by LTSIs (Holly et al., 2019). Our findings thus open exciting new avenues of research on nucleus accumbens LTSIs as a novel target for endogenous and exogenous mu opioid receptor agonists.

## Acknowledgements

Research reported in this publication was supported by the University of Minnesota’s MnDRIVE (Minnesota’s Discovery, Research and Innovation Economy) initiative; the Center for Neural Circuits in Addiction and NIDA Core Center Grant P30DA048742; and National Institutes of Health grants DA037279 and DA048946 (PER). The viral vectors used in this study were generated by the University of Minnesota Viral Vector and Cloning Core. We thank Esther Krook-Magnuson, Emilia Lefevre, Brian Trieu, and all members of the Rothwell lab for helpful discussions.

## Notes

### Competing Interest Statement

The authors have declared no competing interest.

